# Mimicking orchids lure bees from afar with exaggerated ultraviolet signals

**DOI:** 10.1101/2022.07.04.498711

**Authors:** Daniela Scaccabarozzi, Klaus Lunau, Lorenzo Guzzetti, Salvatore Cozzolino, Adrian G. Dyer, Nicola Tommasi, Paolo Biella, Andrea Galimberti, Massimo Labra, Ilaria Bruni, Lorenzo Pecoraro, Giorgio Pattarini, Mark Brundrett, Monica Gagliano

**Author notes:** **Corresponding author:** Daniela Scaccabarozzi **Email:**.

## Abstract

1. Flowers have many sensory traits to appeal to pollinators, including ultraviolet (UV) absorbing markings, which are well known for attracting bees at close proximity (e.g. < 1 m). While striking UV signals have been thought to attract pollinators also at greater distances of meters, how the signals impact the plant pollination success over distance remains unknown. Here we report the case of the Australian orchid *Diuris brumalis,* a non-rewarding species, pollinated by bees via mimicry of rewarding pea plant *Daviesia decurrens.* When distant from the pea plant, *Diuris brumalis* was hypothesized to enhance pollinator attraction by exaggerately mimicking the floral ultraviolet (UV) reflecting patterns of its model.
2. By experimentally modulating floral UV reflectance with a UV screening solution, we quantified the orchid pollination success at variable distance from the model plants.
3. We demonstrate that the deceptive orchid *Diuris brumalis* attracts bee pollinators by emphasizing the visual stimuli, which mimic the floral UV signalling of the rewarding model *D. decurrens.* Moreover, the exaggerated UV reflectance of *D. brumalis* flowers impacted pollinators’ visitation at an optimal distance from *D. decurrens*, and the effect decreased when orchids were too close or too far away from the model.
4. Our findings show that salient UV flower signalling plays a functional role in visual floral mimicry, likely exploiting perceptual gaps in bee neural coding, and mediates the plant pollination success at much greater spatial scales than previously expected.

## 1. Introduction

The art of deception, involving a range of strategies individuals adopt to change the perception and behaviour of others, is commonly practiced by many organisms across the animal and plant kingdoms. Mimicry, a form of deception, allows individuals to conceal their identity and avoid recognition by (more or less) closely imitating the behaviour or resembling the appearance of their models (Dawkins & Krebs, 1979). One of the most remarkable examples of these deceptive adaptations is the duping of pollinating animals by plant mimics. Amongst the 32 families of deceptive plants (Renner, 2006), orchids are undoubtedly the master tricksters. With an estimate of about one-third of all species lacking floral reward to pollinators (Dafni, 1984; Ackerman, 1986a; Jersáková et a.l, 2006), orchids deceive by luring food-seeking animals by fine-tuned mimicry (i.e. Batesian floral mimicry) or general, even inaccurate, resemblance of rewarding flowers (e.g. generalized food deception; Shretha et al., 2020). Surprisingly, how plants succeed in their deception despite widespread imperfect mimicry remains poorly understood (Roy & Widmer, 1999; Schiestl, 2005; Vereecken & Schiestl, 2008). In animals, the success of imperfect mimicry has been explained by high-salience traits, which overshadow other ‘less important’ traits (Cuthill, 2014; Kazemi et al., 2014) by being highly discriminable from the background (Frieman & Reilly, 2015). Although high-salience signals such as attention-grabbing colours and visual patterns occur as frequently in animals (Kazemi et al., 2014) as in plants (Peter & Johnson, 2008; Jersáková et al., 2012; Peter & Johnson, 2013), their role in explaining imperfect mimicry in plants has received comparatively less attention (Vereecken & Schiestl, 2008). In this study, we examined the role salient ultraviolet (UV) signalling plays in the imperfect floral mimicry of a rewardless orchid that falsely advertises a reward to attract bees when afar from model plants.

Flowering plants and pollinating insects interact through a wide range of sensory modalities which affect both the pollinator’s foraging behaviour and the plant’s reproductive success (Leonard et al., 2011a; Glover, 2011). Pollinating insects, in particular bees, make their foraging decisions most effectively by combining visual, olfactory and somatosensory floral signals (Leonard at al., 2011a; Kulahci et al., 2008), yet their innate preference for conspicuous floral displays makes colour and contrasting visual patterns the primary means by which plants first attract them (Naug & Arathi, 2007; van der Kooi et al., 2019). Bees, the main flower visitors, have phylogenetically conserved trichromatic vision (Briscoe & Chittka, 2001) which can be conveniently modelled with maximum sensitivity UV (approx. 340 nm), Blue (435 nm) and Green (560 nm) photoreceptors (Chittka & Kevan, 2005). Plants produce striking floral markings and patterns by absorbing and reflecting UV light (Briscoe & Chittka, 2001; Dinkel & Lunau, 2001; Lunau et al., 2006; Papiorek et al., 2016; Lunau et al., 2021). Interestingly, it is the UV reflectance display rather than UV pattern (absorbance and reflectance) that increases insect visitation (Johnson & Andersson, 2002; Klomberg et al., 2019). The high chromatic contrast that such UV signals can generate is thought to enhance colour salience in an opponent colour system (Lunau et al., 2006; Papiorek et al., 2016; Chittka et al., 2001); however, such chromatic contrast is assumed to work only at relatively short distances of about few centimetres (e.g. UV absorbing “floral guides”; Giurfa et al., 1996; Garcia et al., 2001; Horth et al., 2014; Orbán & Plowright, 2014). This is because bees typically only use the long wavelength green input channel of their visual system to enable fast achromatic processing and detection of small target signals (Klomberg et al., 2019), although some psychophysics shows that alternative chromatic channels may in some cases also be important for bee detection and recognition (Zhang et al., 1995; Morawetz et al., 2013; Dyer et al., 2019). That UV reflectance can also attract pollinator insects from further afield has been posited for decades (Daumer, 1956; Daumer, 1958; Burr et al., 1995; Koski & Ashman, 2014) but remains unverified.

Salient UV signals against the background may be particularly relevant for increasing long distance attractiveness in plants that employ flower mimicry (Dyer, 1996). One such plant is the Australian donkey orchid *Diuris brumalis* whose outer petals appear yellow to human vision, and also reflect large amounts of UV that would be conspicuous to the visual system of bees (Burr et al., 1995). *Diuris brumalis* is a food-deceptive species which secures pollination by resembling the co-occurring rewarding pea plant *Daviesia decurrens* (Scaccabarozzi et al., 2018). The mimicry signals consist of both colour reflectance and inner flower shape, as the outer petals diverge from the pea flower shape (Scaccabarozzi et al., 2018). In addition, the size of the orchid flower is about three times bigger than the pea flower (Fig. 1 a). Whilst the mimicry in size and shape is imperfect, the orchid coloration, with the average colour loci corresponding to the UV region, is perceptually similar to the pea model in colour space; such overlap (< 0.06 colour hexagon units) makes the two species not readily distinguishable in the eyes of their bee pollinator, *Trichocolletes* spp. (Hymenoptera: Collectidae, Fig 1a; Scaccabarozzi et al., 2018). Food-deceptive orchids are known for gaining their pollination success not only by resembling a specific rewarding model flower (Scaccabarozzi et al., 2018; Schaefer & Ruxton, 2009; Dyer et al., 2012), but also exaggerating their floral signals that advertise the false reward and thus increase pollinator responses (Ackerman, 1996b). We hypothesized that the two UV reflecting outer petals of *Diuris* function as exaggerated version (for UV reflectance display) of the floral signal display *Trichocolletes* bees normally encounter in the rewarding *Daviesia* peas. We expected that modulating the *Diuris* exaggerated UV signals over a spatial scale does affect pollination success when orchids are relatively distant from their model food plants because pollinators are more likely to mistake the orchid for the rewarding model when afar. Here we report that the orchid not only uses exaggerated UV reflectance to falsely advertise a potential reward to attract bees from afar, but the ruse works most effectively at an optimal distance of several meters revealing the functional role of salient visual stimuli when mimicry is imperfect.

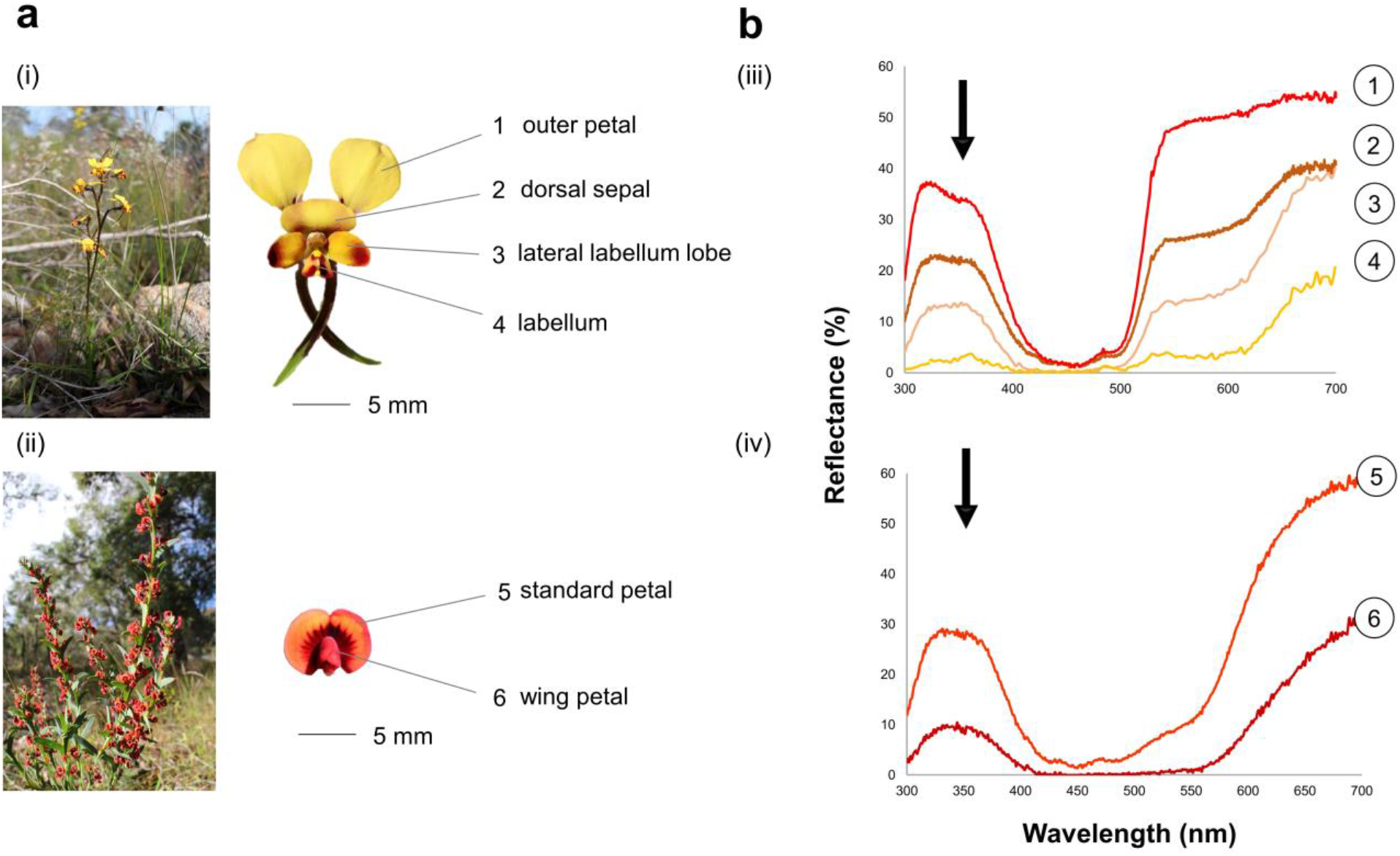

## 2. Materials and Methods

### Study system

Endemic to Western Australia, the orchid *Diuris brumalis* produces yellow–brown nectarless flowers between July and August and is pollinated via mimicry of rewarding pea plants *(Daviesia* spp.) by *Trichocolletes* (Colletideae) native bees (Scaccabarozzi et al., 2018). *Trichocolletes* is a genus of solitary bees that are specialist and speed (visits last less than two seconds) feeder on pea flowers and displaying a distinctive and identical behaviour on both orchids and peas, confirming that it is successfully deceived. The orchid mimics the papilionaceous flower typical of the pea model and while the visible spectrum differs between the mimic and model flower, they are likely to look similar through a bee visual model (Scaccabarozzi et al., 2018). However, the orchid flower diverges from the pea flower structure for exhibiting two prominent outer petals.

We carried out our study in *Diuris brumalis* populations spread along the Darling Range in Western Australia during 2018, 2019 and 2020 (Table S1). *In situ* studies and experimental setting were preferred as the orchids are protected by national regulation and their withdrawl is only allowed for few biological material.

### Floral morphology and colour properties

To test the hypothesis that the two outer petals of *Diuris* may function as an exaggerated version of *Daviesia* floral signals, we determined whether the orchid outer petals had the highest UV reflectance, so amplifying the UV reflectance of the pea model. Firstly, we determined whether the outer petals were the component of the *Diuris* flower with the highest UV spectral reflectance. We obtained UV measurements for each floral component (n = 6 flowers) for both species using a Cary 4000 UV-Vis spectrophotometer (Agilent Technologies, CA) and calculating the average spectral reflectance for each floral part.

Secondly, we measured the size of the flower components of the flower in 10 plants of both *Diuris* and *Daviesia* (Fig. S1; Data source S1). We obtained for both species a UV salient signal ratio according to cut value of Australian flowers following Dyer (1996) (see Data source S1). Flower components’ area were estimated as follows: as flowers of *Diuris* and *Daviesia* show minimal concavity or convexity, the area of the outer and central component of *Diuris* were estimated by approximating the components to the closest geometric figures, the ellipse (orange) and the circle (green), respectively (Fig. S1). *Daviesia* standard petals’ area was approximated to an ellipse, to which was subtracted a secondary minor ellipse circumscribing the wing and keel petals (Fig. S1; Data source S1).

To quantify the contrast of the respective flower signals we used the bee visual parameters according to Chittka and Kevan (2005) and neural coding that enable converting visual signals sensed by each receptor channel into Excitation values between 0 and 1.0. The visual system was adapted to foliage background with a biologically relevant neural resting Excitation value of 0.5 and a contrast of zero (Chittka et al., 1994; Spaethe et al., 2001). This model enables the calculation of absolute contrast values ranging from 0 to 0.5 (maximum contrast) for any stimulus that is different to the background as perceived by the visual system of bees (Table 1).

**Table 1.** Average of excitation values (± SD: standard deviation) of bee photoreceptors (UV, blue, green) according to Chittka^<u>46,47</u>^ and relative corrected values for *Diuris* and *Daviesia* flower components as shown in Fig. 1, including *Diuris* outer petals treated by UV filter. Excitation values range between 0 and 1.0 where a value of 0.5 represents no excitation of the sensory neural channel, and so absolute maximum excitation contrast is 0.5 for each respective channel.

| Flower components | | E(uv)±SD | E(uv)-0.5 | E(b)±SD | E(b)-0.5 | E(g)±SD | E(g)-0.5 |
| --- | --- | --- | --- | --- | --- | --- | --- |
| 1 | *Diuris brumalis* outer petal | 0.84±0.03 | 0.34 | 0.49±0.07 | 0.01 | 0.70±0.03 | 0.20 |
| | *Diuris brumalis* outer petal treated with UV filter | 0.48±0.03 | 0.02 | 0.32±0.07 | 0.18 | 0.70±0.03 | 0.20 |
| 2 | *Diuris brumalis* dorsal sepal | 0.77±0.09 | 0.27 | 0.40±0.09 | 0.10 | 0.57±0.07 | 0.07 |
| 3 | *Diuris brumalis* lateral labellum lobe | 0.64±0.17 | 0.14 | 0.20±0.11 | 0.30 | 0.42±0.17 | 0.08 |
| 4 | *Diuris brumalis* labellum | 0.25±0.17 | 0.25 | 0.07±0.07 | 0.43 | 0.15±0.03 | 0.35 |
| 5 | *Daviesia decurrens* standard petal | 0.77±0.02 | 0.27 | 0.39±0.09 | 0.11 | 0.45±0.06 | 0.05 |
| 6 | *Daviesia decurrens* wing petal | 0.56±0.10 | 0.06 | 0.13±0.05 | 0.37 | 0.14±0.06 | 0.36 |

False colour photography in ‘bee view’ format was used to reveal the overall colour pattern perceived by bees of *Diuris* and *Daviesia* flowers (Fig. 2a, Fig. 2b; see Methods S3 in Supporting Information). Spectrometer measuremens of flower components of *Diuris* and *Daviesia* were converted according to the established bee visual model (Chittka, 1992). Location of colour loci was calculated as from mean of reflectance for floral parts of *Diuris brumalis,* and *Daviesia decurrens* (Fig. 2 c).

**Fig. 2.**
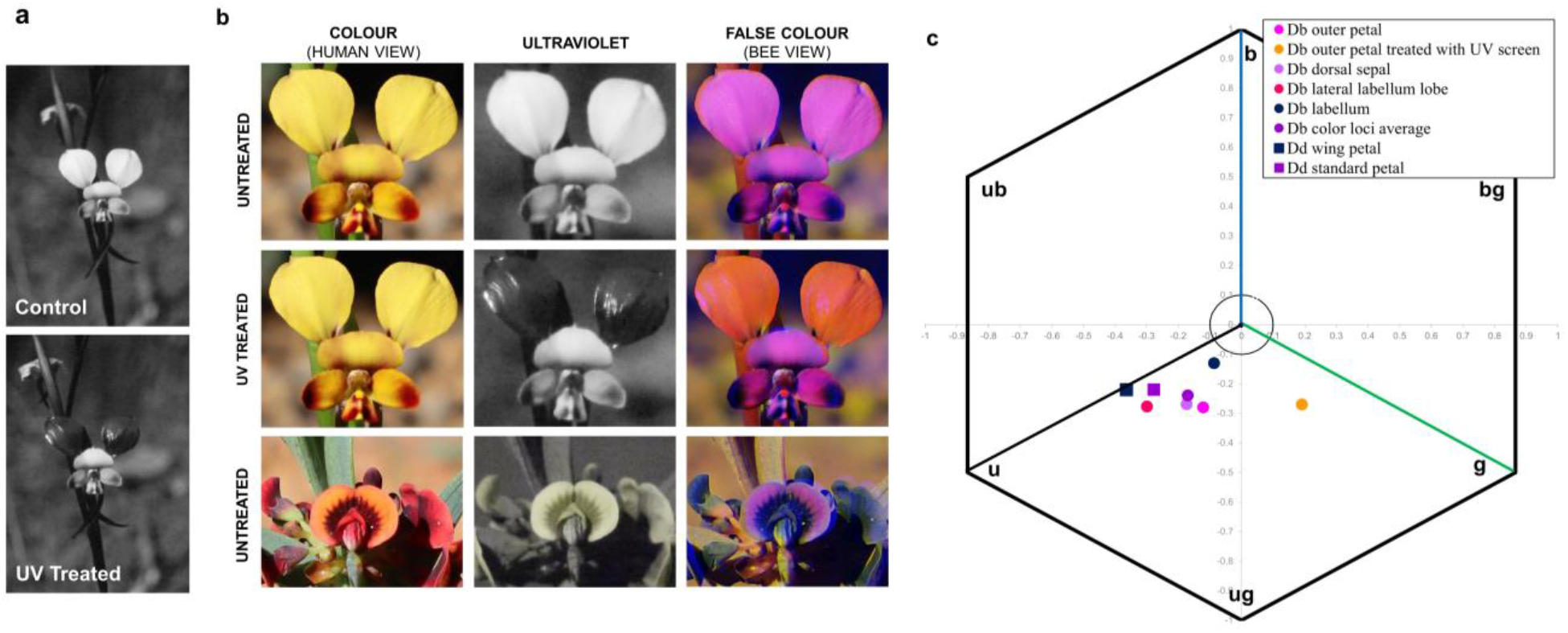
Colour patterns perceived by bees in treated and untreated *Diuris* flowers and untreated *Daviesia*. (**a**) *Diuris* flower photographed in UV before (control, C) and after applying the UV absorbant filter on the outer petals (UV treated, T). (**b**) False colour photography in ‘bee view’ reveals the overall colour pattern perceived by bees in treated (i.e., application of the UV filter solution) and untreated outer petals of *Diuris* flower and untreated *Daviesia* flower. The UV filter is effectively a long pass filter transmitting all wavelengths above 400 nm, free of fragrance, oil, PABA, alcohol, parabens and preservatives (Kinesys, Canada). Importantly, the UV images of treated outer petals show very similar reflectance properties to the background and stem foliage reflectance, confirming that the experimental manipulation knocked out UV signalling with respect to background colouration. (**c**) Location of colour loci was calculated as from mean of reflectance for floral parts of *Diuris brumalis* (Db), and *Daviesia decurrens* (Dd). The calculations were made using the Hexagon colour model of bee vision (Chittka, 1992). This model represents the internal perception of flower colours by bee pollinators, and resultant sectors [u (ultraviolet); ub (ultraviolet-blue); b (blue) bg (blue-green); g (green); ug (ultraviolet-green)] show how bees likely interpret spectral signals].

### Model-mimic distance experiment

To test whether *Diuris* pollination success varies depending on the distance to the model pea plants, in 2019 we first quantified distance between an individual orchid and all the surrounding pea models within a quadrat of 30 x 30 m centred on a single orchid plant (N = 122 orchids across 5 populations; Fig. S2) for all orchid plants per population. As a result, all quadrats were overalapped within the same population, but not among populations (as distance between population was greater than 500 m). To quantify pollination attraction, we recorded the number of pollinaria removed by pollinators in all orchids per population, counting the number of flowers in both orchids and pea plants (pollinia removed in orchids were counted by visually observing the lack of pollinia at the top of the column). We analysed the distance data by using a Generalized Mixed Effect Model (GLMM) with Poisson distribution. The response variable in the model was the number of pollinaria removed and the fixed effects were the distance from the pea model and the number of orchid flowers. Population was treated as random factor, since it was found to be significant in influencing the number of pollinaria removed. The model was evaluated for its dispersion parameter and residuals were evaluated for the assumption of overdispersion and homoschedasticity.

### Ultraviolet manipulations experiments

Subsequent manipulation experiments were carried out in field in 2019 and 2020 by screening the UV properties of the two *Diuris* outer petals with an UV filter solution (see Johnson & Andersson, 2002; Peter & Johnson, 2013), which effectively eliminates UV reflectance whilst transmitting all wavelengths above 400 nm (Fig. 2 a, Fig. 2 b i.e., UV filter). To confirm that treated *Diuris* outer petals did not excite the UV bee photoreceptor as untreated petals and *Daviesia* petals did, we analised the spectral reflectance measurements for the different floral component using the model of bee vision including treated petals (Chittka, 1992; Table 1). False colour photography in ‘bee view’ format was applied on *Diuris* flower with treated outer petals to show the overall colour pattern (Fig. 2 b).

In the first field manipulation experiment (in 2019), we tested the hypothesis that UV reflectance enhances orchid pollination success (pollen removal) only when orchids are out of patch of model pea plants. We quantified the number of pollinaria removed from *Diuris* flowers by free-foraging bees when the mimicking orchid occurred inside [IN] and outside [OUT] the 30 x 30 m patch of model plants (within a maximum distance of 10 meters from the patch; Fig. 3a). The patch size encompassed most orchid plants belonging to an individual population according to former studies on pollination success of *Diuris* at this location (Scaccabarozzi et al., 2018). Over a 4-day period, all orchids in both [IN] and [OUT] groups (N = 400 across 5 populations, Table S1) were treated with the UV filter. Within each group, a randomly selected half of the orchids was sprayed on the front and back of the two outer petals (treatment, T) and the other half of the orchids at the base of the corolla (control, C). Number of flowers was standardised in each group by removing exceeding flowers. The filter was applied before the daily peak of bee activity and left for 3 hours (correspoding to the filter persistence on petals) from 11.00 am to 1.00 pm and the number of pollinia removed from the orchids within each group was recorded during the subsequent two-hour period (1.00 to 3.00 pm). Prior to the filter application, the treated and untreated plants were numbered and tagged. We also recorded the number of pollinia already removed per flower / per plant to make sure of the net counting of pollinia. When revisiting the plants for scoring pollinia, we checked the plants in the same order followed prior to the treatement. Statistics were based on comparisons of removed pollinia between experimental groups (UV-treated petals) and control groups (UV-untreated petals).

**Fig. 3.**
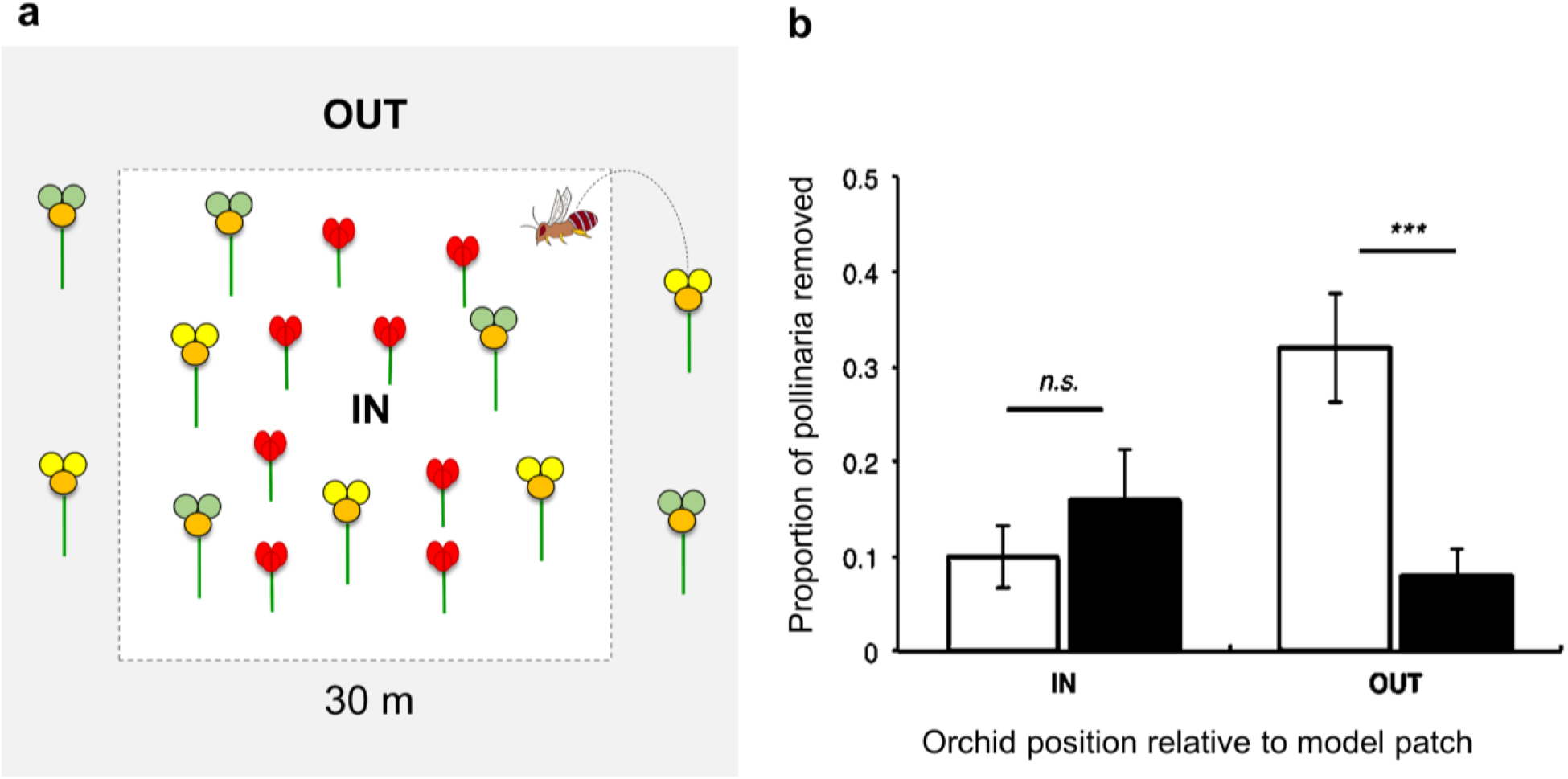
Effect of distance from model plants on *Diuris* pollination success. (**a**) *Diuris* orchids (yellow [untreated] and green [UV treated] flowers) inside [IN] and outside [OUT] a 30 x 30 m patch with *Daviesia* pea (red flowers). (**b**) Mean proportion of pollinaria bees removed from treated (black bars; T) and untreated *Diuris* flowers (white bars; C) relative to the orchid’s distance ([IN] and [OUT]) from the model pea. Each experimental group consists of N = 100 orchids. Error bars are 95% confidence intervals; *n.s.* no significant difference among experimental groups; ***significant difference at Bonferroni corrected α = 0.0125.

In the second field manipulation experiment (in 2020), we tested the hypothesis that by displaying an exaggerated version of *Daviesia’s* attractive UV reflectance, *Diuris* benefits from pollinators that mistake it for the rewarding model from afar. We quantified pollinaria removal within 63 orchid groups randomly selected across three large orchid populations (Table S1, Populations 1,2,3). Each orchid group consisted of two orchid clumps, each containing between 2 and 12 plants. Each orchid clump was selected to be at approximately the same distance from a model pea plant (from 0 to 15 m) at a variable angle from the pea plant (Fig. 4a).

**Fig. 4.**
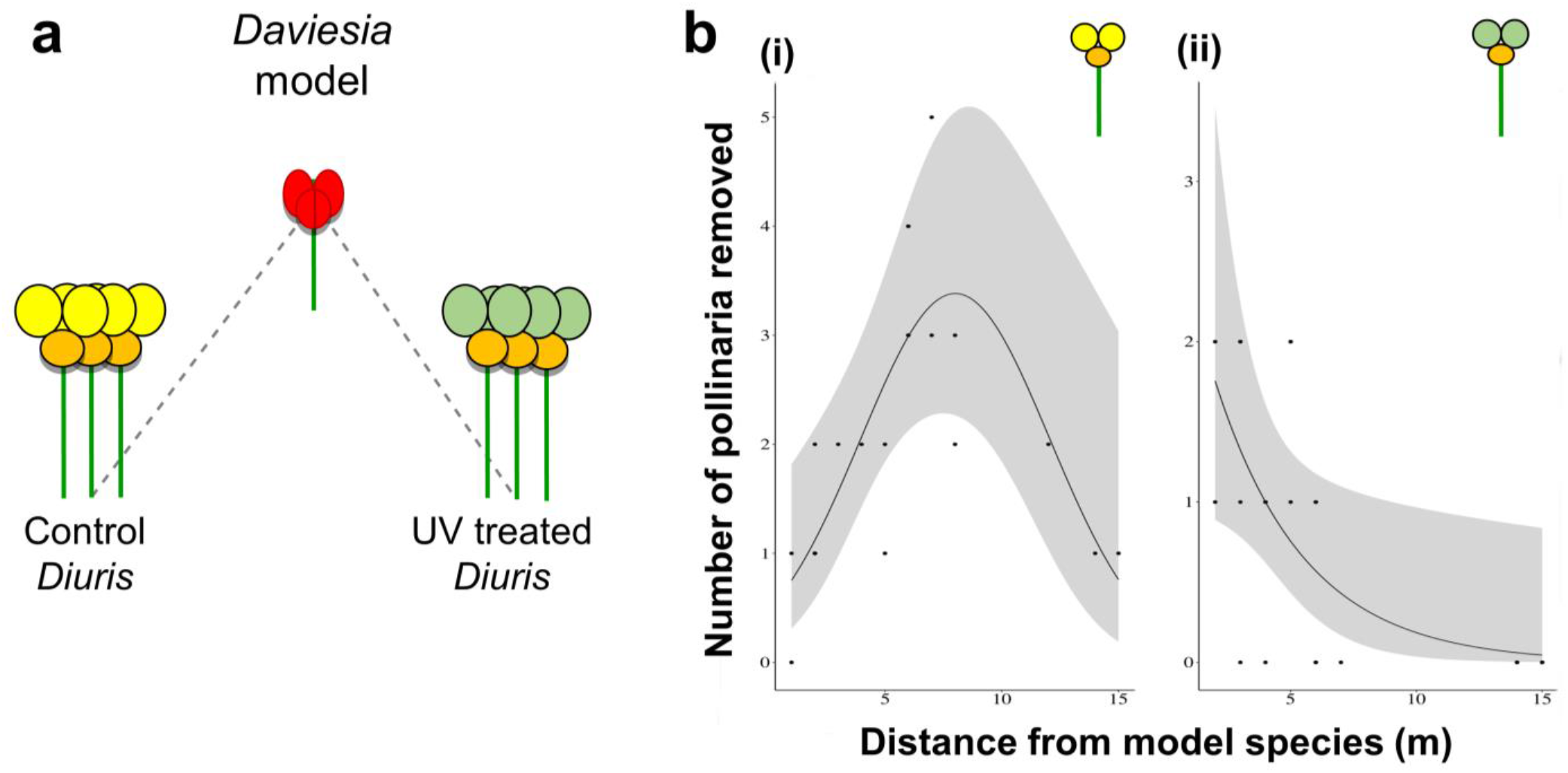
Effect of *Diuris* UV reflectance on the orchid’s pollination success relative to mimic-model distance. (**a**) Experimental set up treated, untreated orchid groups and pea plants, (**b**) Pollinaria removal was quantified in 195 orchids (N = 476 orchid flowers). Pollination success of control *Diuris* relative to distance from *Daviesia* (i) was best described by an inverted parabolic function peaking at ~8 m distance from model pea (χ^2^=9.87, p < 0.05 for the squared and linear term respectively) (N=238 flowers, n =43 pollinia removed). Pollination success of UV treated orchids (ii) exhibited an exponential decrease with distance from model pea plants (χ^2^=10.26, p < 0.001); (N=238 orchid flowers, n =17 pollinia removed). Refer to Data source S7 for full data.

Within each orchid group, *Diuris* floral display (i.e., number of flowers in each clump) was standardized by removing excess flowers. Within each group, the UV filter solution was sprayed on the outer petals of one clump (treatment, T) and at the base of the corolla on the other clump (control, C) as in the previous experiment (same treatment and plant visitation timing). Prior to the UV filter application, the treated and untreated plants were numbered, tagged and the number of pollinia removed per flower / per plant was recorded. Pea plant flowers range was uniform among plants at the time of experiment. The number of pollinaria removed from the UV treated and control orchids within each group was recorded as a function of of the orchid’s distance to the pea plant and was modelled by a Poisson GLMM (appropriate for count data) with a fixed effect for treatment.

## 3. Results

### Outer petals in the mimic orchid display an exaggerated UV signal of model plants

Spectral reflectance and morphological measurements of flowers components confirmed that *Diuris* functioned as an exaggerated version of the floral signals bees normally encounter in the rewarding *Daviesia* peas. The outer petals proved to be both the largest component of the orchid flower and the area with the highest UV reflectance (Fig. 1b; Fig S1, Data source S1). *Diuris* flowers produced an overall higher UV salient signal ratio (ratio=0.84) comparing to *Daviesia* flowers (ratio=0.69; Data source S1) representing a 22% increase of surface area reflecting UV signal (Data source S1). The strength of the UV signalling in *Diuris* had a contrast value of 0.34 which is 26% greater than the UV channel contrast value of 0.27 in *Daviesia* standard petals (Table 1). False colour photography in ‘bee view’ revealed the similarity of the overall colour pattern perceived by bees of *Diuris* and *Daviesia* flowers (Fig. 2 b).

According to the colour model, petals of *Diuris* and petals of *Daviesia* are located in the bee perceived “ug” (UV-green) and “u” (ultraviolet) sectors of the Hexagon colour space related to the excitation of bee photoreceptors and subsequent bee neural coding of information (Fig. 2 c; Table 1) (Chittka, 1992; Chittka et al., 1994).

### Orchid pollination success relate to mimic-model distance

Mimic-model distance on large scale revealed that the number of pollinaria removed from the orchid flowers decreased significantly with distance between orchid and pea (Fig. S2). Specifically, pollination success decreased significantly with the square of orchids’ distance from the pea model (χ^2^=17.09, p < 0.001) and was compromised when the distance between orchid and pea was greater than 15 m (χ^2^=9.49, p = 0.002).

### The exaggerated UV signal enhances the orchid success relative to model plants

Field manipulation experiments showed that the exaggerated UV signal of *Diuris* outer petals enhances the orchid’s pollination success. The UV filter treatment had no attracting or repelling effect on the pollinators (see Methods S1 in Supporting Information).

Treated petals of *Diuris brumalis* are located in the bee perceived “g” (green) Hexagon sector and did not excite bee UV photoreceptors (Fig. 2 c; Table 1). Secondly, the colour model confirmed that excitation of Green receptor, which is known to be important for how bees efficiently find flowers (Giurfa et al., 1996; Skorupski & Chittka, 2010; Garcia et al., 2021), was not affected by UV filter treatment (Table 1). False colour photography in ‘bee view’ confirmed that the UV filter knocked out UV signalling with respect to background colouration (Fig. 2 b).

In the first field manipulation experiment, we quantified the number of pollinaria removed from treated and control *Diuris* flowers by free-foraging bees when the mimicking orchid co-occurred with the model pea within a 30 x 30-m patch per orchid population [IN] and when the mimics occurred outside the patch of model plants [OUT] (Fig. 3a). The application of the UV filter on the two outer petals resulted in a significant effect on the number of pollinaria removed by bees from the orchid flowers (χ^2^=19.81, p < 0.001). There was no difference in the pollination success of *Diuris* whose outer petals had been treated with the UV filter [IN-T] compared to untreated control orchids [IN-C] inside the patchs of model plants (Fig. 3b). Outside the patchs of model plants, however, orchids with UV-filter treatment [OUT-T] experienced significantly lower pollinaria removal than control ones [OUT-C] (Fig. 3b).

In the second field manipulation experiment, we found that pollination success of control *Diuris* increased with distance by peaking at ~8 m away from the model peas before declining and becoming ineffectual at distances greater that 15 m (Fig. 4bi). We detected no effect of UV reflectance on *Diuris* pollination success when the orchids were closer than a few meters to their model pea plants (Fig. 4bi, Fig. 4bii).

## 4. Discussion

Our results establish that *Diuris* orchids mimic and exaggerate *Daviesia*’s attractive floral signals in terms of UV reflectance, display, and contrast as perceived by bee pollinators. Flowers that reflect greater than 10% UV radiation, like *Diuris* and *Daviesia,* are shown to have evolved this salient trait to improve communication with bees since most organic background material like leaf foliage has very low UV reflectance (Dyer, 1993; Chittka et al., 1994; Spaethe et al., 2001; van der Kooi et al., 2019).

By masking the UV reflectance in half of the orchids inside the *Daviesia’s* patch, the treatment effectively made those *Diuris* displaying the exaggerated UV signal a rarer phenotype, which would be predicted to enjoy greater pollination success by negative frequency-dependent selection (Schiestl, 2005; Schiestl & Johnson, 2013). Instead, there was no difference in the pollination success of *Diuris* whose outer petals had been UV screened [IN-T] compared to untreated control orchids [IN-C] inside the *Daviesia’s* patch (Fig. 3b). At closer range, within pea patch, bees apparently recognise plants by spotting other visual traits as the shape of *Diuris* two outer petals. For example, a colour trait may become less effective in ensuring successful mimicry when other secondary traits such as size and shape of the flowers can be better discriminated (Gigord et al., 2002; Johnson et al., 2006). Outside the model patch, however, orchids with UV-filter treatment [OUT-T] experienced substantially lower pollinaria removal than control ones [OUT-C] (Fig. 3b), due to a lack of the salient signal which is associated to the model trait. Thus, the exaggerated UV signal produced by *Diuris* outer petals only increased the orchid’s pollination success when the mimic was further away from its models’ patch. Our findings demonstrate that salient floral UV reflectance plays a critical role in ensuring *Diuris* pollination success and explain why the exaggerated UV signal is strategically relevant in floral mimicry when the model is not very close to the mimic. According to previous theories predicting the effectiveness of the mimic’s floral stimuli to decline with distance from its model (Johnson & Schiestl, 2016; Duffy & Johnson, 2017), we also found that the number of pollinaria removed from the orchid flowers decreased significantly with distance between orchid and pea (Fig. S2). However, the strength and direction of this effect may vary across different spatial scales and conclusions about the importance of floral stimuli will depend on the scales at which studies are undertaken. For example, by examining the mimic-model effect at considerably smaller spatial scales than usually investigated (i.e., tens to hundreds of meters) (Duffy & Johnson, 2017; Johnson et al., 2003; Peter & Johnson, 2008), our results show that the exaggerated UV reflectance of *Diuris* outer petals function to enhance pollination at an optimal model-mimic range of ~8 m. *Diuri*s outer petals might promote pollinator deception via bee cognitive misclassification (Dyer et al, 2012; Johnson & Schiestl, 2016), displaying colour frequencies below the optimal range of colour disciminations in hymenopteran (i.e. 400-500 nm) (Peitsch et al., 1992), especially for free-flying honeybees (von Helversen, 1972; Rohde et al., 2013).

But why might the observed distance range from model species be optimal? To understand this question, we must delve into both the neurophysiology and physiology of how bee pollinators perceive their world. When a bee receives sweet tasting nectar reward from a rewarding plant like *Daviesia decurrens,* this promotes a sustained positive neural response via the ventral unpaired median (VUM) neurons that permit an association between flower and reward with a sustained spiking response of about 15s (Hammer, 1993; Perry, 2013), and can enable simple associative learning of colour information (Dyer & Chittka, 2004; Giurfa, 2004). It is also known that precise colour memory in both bees and humans requires simultaneous viewing conditions that decay in less than a second once a target model is no longer in view (Uchikawa & Ikeda, 1981; Dyer & Neumeyer, 2005); therefore, being close to a model species might allow a bee to identify potential differences that unmask the deception (von Helversen, 1972). Given that bees may fly up to about 7 m in a second (Spaethe et al., 2001; Srinivasan & Lehrer, 1985), we hypothesize the 8 m distance we observed for optimal pollination success is beyond the theoretical upper limit where precise colour vision operates; at such distances, the bee has to recall from memory what it thought was rewarding and tends to prefer a slightly more salient comparative stimulus, an effect related to peak shift discrimination (Lynn et al., 2005; Leonard et al., 2011b; Martínez-Harms et al., 2014). The fast visits of *Trichocolletes* bees on both model and mimic flowers (Scaccabarozzi et al., 2018), suggest that *Diuris* benefits from foraging speed behaviour that unfavours the accuracy of bee choices (Chittka et al., 2003). Thus, we propose that orchids like *Diuris* master deception by employing both exaggerated signalling and by exploiting the perceptual gaps in pollinators’ visual processing.

Our results also highlight that we gain a very different understanding of the relative role of floral signals if we work at one scale over another and consider the dynamics of pollinator perception. For example, orchid pollination success was greatest when the mimics where further away from their models (e.g. ouside the patch), but within a maximum distance of 10 meters from the model patch. Because the pollination success of deceptive species can be subject to both competition and facilitation effects depending on the density of rewarding (Julliet et al., 2007) and conspecific plants (Duffy & Stout, 2011) the competition orchids experienced within the patch of floriferous pea plants would have been at its strongest (Fig. 3b). However when we accounted for both floral density of conspecific and model plants along a continuous and wider spatial scale (Fig. S2), the pollination success pronouncedly declined at distances greater than 15 m from model plants. At such distances, the orchids no longer had to contend with the peas for pollinators’ attention but the beneficial effect of facilitation between the the plant species also disapperared. Therefore, the importance of exaggerated UV reflectance in attracting pollinators from a range of several meters can be missed and/or mistakenly dismissed if not measured at the scale at which it has its strongest ecologically-relevant effect. Such a long-range signal might not be suspected considering the typical acuity range of bee chromatic vision for stationary stimuli within the confined space of a Y-maze (Giurfa et al., 1996). Overall, our results demonstrate that the functional role of UV reflectance signalling is contingent on the relative distance between deceptive and rewarding species and their pollinators; the distance described here operates at spatial scales of meters, which are much greater than previously described for floral colours. The terminal position of the outer petals on a long stemmed plant (Fig. 1a) likely promotes (wind) movement of this exaggerated UV signal that can be even better perceived from afar by foraging bees (Stojecev et al, 2011; Brock et al., 2016) by acting as a ‘flag signal’. Contributing to a range of floral displays aimed at pollinator senses, UV reflectance acts as an important visual cue in many flowering plant species (Johnson & Andrersson, 2002; Klomberg et al., 2019). The high UV reflectance of *Diuris* outer petals enables bees to find these relatively scarce flowers from a distance of meters. Selection may favour deceptive floral displays capable of longer range UV signalling that help pollinators such as solitary bees to locate flowers in habitats where the distribution of rewarding model flowers is patchy, explaining why relatively large, salient UV signals with high background contrast have evolved in the mimic (Rohde et al., 2013). By revealing that floral salient UV displays are efficiently used by bees not only at the very close ranges already well-documented, but also from further afield (see also Supporting information, Methods S2), we may explain how plant deception succeeds despite imperfect floral mimicry. This finding invite us to extend our understanding of the adaptive significance of UV reflectance and salient signalling that plants display in an captivating phenomenon such as the floral mimicry and more general in nature.

## Acknowledgments

We thank A. Aromatisi for fieldwork assistance and useful discussions on the study design. We acknowledge C. Best for fieldwork assistance, T. Houston for input in the behavioural ecology of native bees, T. Scalzo, P. Chapman, M. Massi and C. May for technical and laboratory assistance.

We acknowledge the following funding sources: Endeavour Fellowship Program grant ID 5117_2016 (DS); Australian Orchid Foundation grant 308.16 (DS); Curtin University grant CIPRS-CSIRS_2017 (DS); Università degli Studi di Napoli Federico II, Short mobility program D.M. 976_2017 (DS); Templeton World Charity Foundation grant TWCF0541 (MG); Australian Research Council Discovery Project DP160100161 (AGD).

## Author contributions

DS, SC, MG, KL, AGD conceived the ideas and designed methodology; DS, MB, AG collected the data; LG, DS, AG, SC, AGD analysed the data; DS, MG, KL, SC led the writing of the manuscript. All authors contributed critically to the drafts and gave final approval for publication.

## Data availability statement

Data needed to evaluate the conclusions in the paper are presented in the Supplementary Information.

## References

Ackerman, J. D. (1986a). Mechanisms and evolution of food-deceptive pollination systems in orchids. Lindleyana, 1, 108–113. https://doi.org/10.1017/S1464793105006986

Ackerman, J. D. (1986b). Coping with the epiphytic existence: pollination strate gies. Selbyana, 9, 52–60.

Briscoe, A., & Chittka, L. (2001). The evolution of colour vision in insects. Annual Review of Entomology, 46, 471–510. https://doi.org/10.1146/annurev.ento.46.1.471

Brock, M. T., Lucas, L. K., Anderson, N. A., Rubin, M. J., Cody Markelz, R. J., Covington, M. F., Devisetty, U. K., Chapple, C., Maloof, J. N., & Weinig, C. (2016). Genetic architecture, biochemical underpinnings and ecological impact of floral UV patterning. Molecular Ecology, 25, 11,22–1140. https://doi.org/10.1111/mec.13542

Burr, B., Rosen, D., & Barthlott, W. (1995). Untersuchungen zur Ultraviolettreflexion von Angiospermenblüten III. Dilleniidae und Asteridae. Tropische und subtropische Pflanzenwelt, 93, 1–175.

Chittka, L. 1992. The colour hexagon: a chromaticity diagram based on photoreceptor excitations as a generalized representation of colour opponency. The Journal of Comparative Physiology A, 170, 533–543. https://doi.org/10.1007/BF00199331

Chittka, L., Shmida, A. V. I., Troje, N., & Menzel, R. (1994). Ultraviolet as a component of flower reflections, and the colour perception of Hymenoptera. Vision Research. 34(11), 1489–1508. https://doi.org/10.1016/0042-6989(94)90151-1

Chittka, L., Spaethe, J., Schmidt, A., & Hickelsberger, A. (2001). Adaptation, constraint, and chance in the evolution of flower colour and pollinator colour vision in Cognitive Ecology of Pollination, L. Chittka, J. D. Thomson, Eds. Cambridge, UK: Cambridge Univ. Press. https://doi.org/10.1017/CBO9780511542268.007

Chittka, L., Dyer, A. G., Bock, F., & Dornhaus A. (2003). Bees trade off foraging speed for accuracy. Nature 424(6947), 388–388. https://doi.org/10.1038/424388a

Chittka, L., & Kevan, P. G. (2005). Flower colour as advertisement in Practical Pollination Biology, A. Dafni, P. G. Kevan, B.C. Husband, Eds. Cambridge, ON, Canada: Enviroquest Ltd.

Cuthill, I. C. (2014). Evolution: the mystery of imperfect mimicry. Current Biology, 24(9), 364–366. https://doi.org/10.1016/j.cub.2014.04.006

Dafni, A. (1984). Mimicry and deception in pollination. The Annual Review of Ecology, Evolution, and Systematics, 15, 259–278. https://doi.org/10.1146/annurev.es.15.110184.001355

Daumer, K. (1956). Reizmetrische Untersuchung des Farbensehens der Bienen. Zeitschrift fur vergleichende Physiologie, 38, 413–478. https://doi.org/10.1007/BF00340456

Daumer, K. (1958). Blumenfarben, wie sie die Bienen sehen. Zeitschrift fur vergleichende Physiologie, 41, 49–110. https://doi.org/10.1007/BF00340242

Dawkins, R., & Krebs, J. R. (1979). Arms races between and within species. Proceedings of the Royal Society B: Biological Sciences, 205, 489–511. https://doi.org/10.1098/rspb.1979.0081

Dinkel, T. & Lunau, K. (2001). How drone flies *(Eristalis tenax* L., Syrphidae, Diptera) use floral guides to locate food sources. Journal of Insect Physiology, 47, 1111–1118. https://doi.org/10.1016/S0022-1910(01)00080-4

Duffy, K. J. & Stout, J. C. (2011). Effects of conspecific and heterospecific floral density on the pollination of two related rewarding orchids. Plant Ecology 212(8), 1397–1406. https://doi.org/10.1007/s11258-011-9915-1

Duffy, K. J. & Johnson, S. D. (2017). Effects of distance from models on the fitness of floral mimics. Plant Biology, 19, 438–443. https://doi.org/10.1111/plb.12555

Dyer, A. G. (1996). Reflection of near-ultraviolet radiation from flowers of Australian native plants. Australian Journal of Botany, 44, 473–488. https://doi.org/10.1071/BT9960473

Dyer, A. G. & Chittka, L. (2004). Fine colour discrimination requires differential conditioning in bumblebees. Die Naturwissenschaften, 91, 224–227. https://doi.org/10.1007/s00114-004-0508-x

Dyer, A. G., & Neumeyer, C. (2005). Simultaneous and successive colour discrimination in the honeybee *(Apis mellifera)*. Journal of Comparative Physiology A, 191, 547–557. https://doi.org/10.1007/s00359-005-0622-z

Dyer, A.G., Boyd-Gerny, S., McLoughlin, S., Rosa, M.G.P., Simonov, V., & Wong, B.B.M. (2012). Parallel evolution of angiosperm colour signals: common evolutionary pressures linked to hymenopteran vision. Proceedings of the Royal Society B: Biological Sciences, 279, 3606–3615. https://doi.org/10.1007/s00359-016-1101-4

Dyer, A.G., Boyd-Gerny, S., Shrestha, M., Garcia, J. E., van der Kooi, C., & Wong, B.B.M. (2019). Colour preferences of *Tetragonula carbonaria* Sm. stingless bees for colour morphs of the Australian native orchid *Caladenia carnea*. Journal of Comparative Physiology A, 205, 347–361. https://doi.org/10.1007/s00359-016-1101-4

Frieman, J. & Reilly, S. (2015). Learning: A behavioral, cognitive, and evolutionary synthesis. Thousand Oaks, CA: Sage Publications.

Garcia, J.E., Dyer, A.G., Burd, M., & Shrestha, M. (2001). Flower colour and size signals differ depending on geographical location and altitude region. Plant Biology, 23, 905–914. https://doi.org/10.1111/plb.13326

Gigord, L. D, Macnair, M. R., Stritesky, M., & Smithson, A. (2002). The potential for floral mimicry in rewardless orchids: an experimental study. Proceedings of the Royal Society B: Biological Sciences 269(1498), 1389–1395. https://doi.org/10.1098/rspb.2002.2018

Giurfa, M., Vorobyev, M., Kevan, P., & Menzel, R. (1996). Detection of coloured stimuli by honeybees: minimum visual angles and receptor specific contrasts. Journal of Comparative Physiology A, 178, 699–709. https://doi.org/10.1007/BF00227381

Giurfa, M. (2004). Conditioning procedure and colour discrimination in the honeybee *Apis mellifera*. Die Naturwissenschaften, 91, 228–231. https://doi.org/10.1007/s00114-004-0530-z

Glover, B. J. (2011). Pollinator attraction: The importance of looking good and smelling nice. Current Biology, 21, R307–R309. https://doi.org/10.1016/j.cub.2011.03.061

Hammer, M. (1993). An identified neuron mediates the unconditioned stimulus in associative olfactory learning in honeybees. Nature, 366, 59–63. https://doi.org/10.1038/366059a0

von Helversen, O. (1972). Information Processing in the Visual Systems of Anthropods, Symposium Held at the Department of Zoology, University of Zurich, R. Wehner. New York, US: Springer Publishing.

Horth, L., Campbell, L., and Bray, R. (2014). Wild bees preferentially visit Rudbeckia flower heads with exaggerated ultraviolet absorbing floral guides. Biology Open, 3, 221–230. https://doi.org/10.1242/bio.20146445

Jersáková, J., Johnson, S. D., & Kindlmann, P. (2006). Mechanisms and evolution of deceptive pollination in orchids. Biological Reviews, 81(2), 219–235. https://doi.org/10.1017/S1464793105006986

Jersáková, J., Jürgens, A., Šmilauer, P., & Johnson S. D. (2012). The evolution of floral mimicry: identifying traits that visually attract pollinators. Functional. Ecology, 26(6), 1381–1389. https://doi.org/10.1111/j.1365-2435.2012.02059.x

Johnson, S. D., & Andersson, S. (2002). A simple field method for manipulating ultraviolet reflectance of flowers. Canadian Journal of Botany, 80, 1325–1328. https://doi.org/10.1139/b02-116

Johnson, S. D., Peter, C. I., Nilsson, L., & Agren, A.J. (2003). Pollination success in a deceptive orchid is enhanced by co-occurring rewarding magnet plants. Ecology, 84, 2919–2927. https://doi.org/10.1890/02-0471

Johnson, S. D., & Morita, S. (2006). Lying to Pinocchio: floral deception in an orchid pollinated by long-proboscid flies. Botanical Journal of the Linnean Society, 152(3), 271–278. https://doi.org/10.1111/j.1095-8339.2006.00571.x

Johnson, S. D., & Schiestl, F. P. (2016). Floral mimicr. Oxford, UK: Oxford University Press.

Julliet, N., Gonzales, M. A., Page, P. A., & Gigord, L. D. B. (2007). Pollination of the European food-deceptive *Traunsteinera globosa* (Orchidaceae): the importance of nectar-producing neighbouring plants. Plant Systematics and Evolution, 265, 123–129. https://doi.org/10.1007/s00606-006-0507-9

Johnson, S. D., Morita, S. Lying to Pinocchio: floral deception in an orchid pollinated by long-proboscid flies. Botanical Journal of the Linnean Society, 152(3), 271–278 (2006). https://doi.org/10.1111/j.1095-8339.2006.00571.x

Kazemi, B., Gamberale-Stille, G., Tullberg, B. S., & Leimar, O. (2014). Stimulus salience as an explanation for imperfect mimicry. Current Biology, 24(9), 965–969. https://doi.org/10.1016/j.cub.2014.02.061

Klomberg, Y., Dywou Kouede, R., Bartoš, M., Mertens, J. E., Tropek, R., Fokam, E. B., & Janeček, Š. (2019). The role of ultraviolet reflectance and pattern in the pollination system of *Hypoxis camerooniana* (Hypoxidaceae). AoB Plants, 11(5), plz057. https://doi.org/10.1093/aobpla/plz057

van Der Kooi, C.J., Dyer, A.G., Kevan, P.G., & Lunau, K. (2019). Functional significance of the optical properties of flowers for visual signalling. Annals of Botany, 123, 263–276. https://doi.org/10.1093/aob/mcy119

Koski, M. H., & Ashman, T. L. (2014). Dissecting pollinator responses to a ubiquitous ultraviolet floral pattern in the wild. Functional Ecology, 28, 868–877. https://doi.org/10.1111/1365-2435.12242

Kulahci, I. G., Dornhaus, A., & Papaj D. R. (2008). Multimodal signals enhance decision making in foraging bumble-bees. Proceedings: Biological Sciences, 275, 797–802. http://www.jstor.org/stable/25249577

Leonard, A. S., Dornhaus, A., & Papaj, D. R. Forget-me-not: Complex floral displays, inter-signal interactions, and pollinator cognition. Current Zoology, 57, 215–224 (2011a). https://doi.org/10.1098/rspb.2007.1176

Leonard, A. S., Dornhaus, A., & Papaj, D. R. Flowers help bees cope with uncertainty: signal detection and the function of floral complexity. Journal of Experimental Biology, 214, 113–121 (2011b). https://doi.org/10.1098/rspb.2007.1176

Lunau, K., Fieselmann, G., Heuschen, B., & Van De Loo A. (2006). Visual targeting of components of floral colour patterns in flower-naïve bumblebees (*Bombus terrestris*; Apidae). Naturwissenschaften 93, 325–328. https://doi.org/10.1007/s00114-006-0105-2

Lunau, K., Scaccabarozzi, D., Willing, L., & Dixon, K.W. (2021). A bee’s eye view of remarkable floral colour patterns in the Southwest Australian biodiversity hotspot revealed by false colour photography. Annals of Botany, 128, 821–834. https://doi.org/10.1093/aob/mcab088

Lynn, S. K., Cnaani, J., Papaj, D. R., & Björklund, M. (2005). Peak shift discrimination learning as a mechanism of signal evolution. Evolution, 59, 1300–1305. https://doi.org/10.1111/j.0014-3820.2005.tb01780.x

Martínez-Harms, J., Márque, N., Menzel, R., & Vorobyev, M. (2014). Visual generalization in honeybees: evidence of peak shift in colour discrimination. Journal of Comparative Physiology A, 200, 317–325. https://doi.org/10.1007/s00359-014-0887-1

Morawetz, L., Svoboda, A., Spaethe, J., & Dyer, A.G. (2013). Blue colour preference in honeybees distracts visual attention for learning closed shapes. Journal of Comparative Physiology A, 199, 817–827. https://doi.org/10.1007/s00359-013-0843-5

Naug, D., & Arathi, H.S. (2007). Receiver bias for exaggerated signals in honeybees and its implications for the evolution of floral displays. Biology Letters, 3, 635–637. https://doi.org/10.1098/rsbl.2007.0436

Orbán, L. L., & Plowright, C.M.S. (2014). Getting to the start line: how bumblebees and honeybees are visually guided towards their first floral contact. Insectes Sociaux, 61, 325–336. https://doi.org/10.1007/s00040-014-0366-2

Papiorek, S., Junker, R.R., Alves-dos-Santos, I., Melo, G.A.R., Amaral-Neto, L.P., Sazima, M., Wolowski, M., Freitas, L., & Lunau K. (2016). Bees, birds and yellow flowers: Pollinator-dependent convergent evolution of UV patterns. Plant Biology, 18, 46–55. https://doi.org/10.1111/plb.12322

Peitsch, D., Fietz, A., Hertel, H., de Souza, J., Ventura, D.F., & Menzel, R. (1992). The spectral input systems of hymenopteran insects and their receptor-based colour vision. Journal of Comparative Physiology A, 170, 23–40. https://doi.org/10.1007/BF00190398

Perry, C. J., & Barron, A. B. (2013). Neural mechanisms of rereferences ward in insects. Annual Review of Entomology, 58, 543–562. https://doi.org/10.3758/s13415-020-00842-0

Peter, C. I., & Johnson, S. D. (2008). Mimics and magnets: the importance of color and ecological facilitation in floral deception. Ecology, 89(6), 1583–1595. http://hdl.handle.net/10962/d1005977

Peter, C. I., & Johnson, S. D. (2013). Generalized food deception: colour signals and efficient pollen transfer in bee-pollinated species of *Eulophia* (Orchidaceae). Botanical Journal of the Linnean Society, 171(4), 713–729. https://doi.org/10.1111/boj.12028

Renner, S. S. (2006). Rewardless flowers in the angiosperms and the role of insect cognition in their evolution in Plant-pollinator interactions: from specialization to generalization. Oxford, UK: Oxford University Press, pp. 123–144.

Rohde, K., Papiorek, S., & Lunau, K. (2013). Bumblebees *(Bombus terrestris)* and honeybees *(Apis mellifera)* prefer similar colours of higher spectral purity over trained colours. Journal of Comparative Physiology A, 199, 197–210. https://doi.org/10.1007/s00359-012-0783-5

Roy, B. & Widmer, A. (1999). Floral mimicry: A fascinating yet poorly understood phenomenon. Trends Plant Science, 4, 325–330.

Scaccabarozzi, D., Cozzolino, S., Guzzetti, L., Galimberti, A., Milne, L., Dixon, K. W., & Phillips, R.D. (2018). Masquerading as pea plants: behavioural and morphological evidence for mimicry of multiple models in an Australian orchid. Annals of Botany, 122, 1061–1073. https://doi.org/10.1093/aob/mcy166

Schaefer, H.M., & Ruxton, G.D. (2009). Deception in plants: mimicry or perceptual exploitation? Trends in Ecology & Evolution, 24, 676–685. https://doi.org/10.1016/j.tree.2009.06.006

F. P. Schiestl, On the success of a swindle: Pollination by deception in orchids. (2005). Naturwissenschaften, 92, 255–264. https://doi.org/10.1007/s00114-005-0636-y

Schiestl, F.P., & Johnson, S.D. (2013). Pollinator-mediated evolution of floral signals. Trends in Ecology & Evolution, 28, 307–315. https://doi.org/10.1016/j.tree.2013.01.019

Shrestha, M., Dyer, A. G., Dorin, A., Ren, Z. X., & Burd, M. (2020). Rewardlessness in orchids: how frequent and how rewardless?. Plant Biology, 22(4), 555–561. https://doi.org/10.1111/plb.13113

Skorupski P., & Chittka L. (2010). Differences in photoreceptor processing speed for chromatic and achromatic vision in the bumblebee, *Bombus terrestris*. The Journal of Neuroscience, 30, 3896–3903. https://doi.org/10.1523/JNEUROSCI.5700-09.2010

Spaethe, J., Tautz, J., & Chittka, L. (2001). Visual constraints in foraging bumble bees: flower size and colour affect search time and flight behavior. Proceedings of the National Academy of Sciences of the United States of America, 98, 3898–3903. https://doi.org/10.1073/pnas.071053098

Srinivasan, M. & Lehrer, M. (1985). Temporal resolution of colour vision in the honeybee Journal of Comparative Physiology A, 157(5), 579–586. https://doi.org/10.1007/BF01351352

Stojcev, M., Radtke, N., D’Amaro, D., Dyer, A.G. & Neumeyer, C. (2011). General principles in motion vision: colour-blindness of object motion depends on pattern velocity in honeybee and goldfish. Visual Neuroscience, 28, 361–370. https://doi.org/10.1017/S0952523811000101

Uchikawa K., & Ikeda, M. (1981). Temporal deterioration of wavelength discrimination with successive comparison method. Vision Research, 21, 591–595. https://doi.org/10.1016/0042-6989(81)90106-1

Vereecken, N. J. & Schiestl, F. P. (2008). The evolution of imperfect floral mimicry. Proceedings of the National Academy of Sciences of the United States of America, 105(21), 7484–7488. https://doi.org/10.1073/pnas.0800194105

Zhang, S.W., Srinivasan, M.V., & Collett, T. (1995). Convergent processing in honeybee vision: multiple channels for the recognition of shape. Proceedings of the National Academy of Sciences of the United States of America, 92, 3029–3031. https://doi.org/10.1073/pnas.92.7.3029

