## Supporting information for "Mimicking orchids lure bees from afar with exaggerated ultraviolet signals"

***Corresponding author:** Daniela Scaccabarozzi


**This file includes:**

Figures and legends S1 to S3

Supplementary text

Table S1

SI References

Legends for Datasets S1 to S7

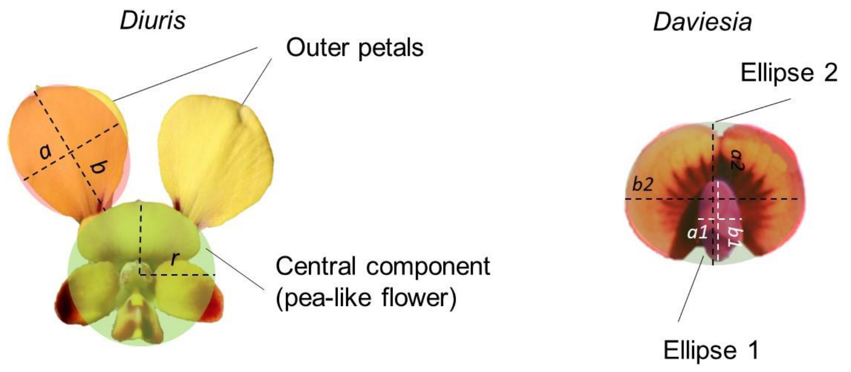


**Fig. S1.** Morphological components of *Diuris* and *Daviesia* flowers. The height (a) and width (b) of *Diuris* outer petals forming an approximated ellipse and the radius (r) of its central component (forming an approximated circle) comprised of dorsal sepal plus labellum and labellum lobes. The width (a1) height (b1) of the wing petals comprising the keel edge and the height (a2) width (b2) of

*Daviesia* standard petals formed approximated ellipses 1 and 2, respectively.

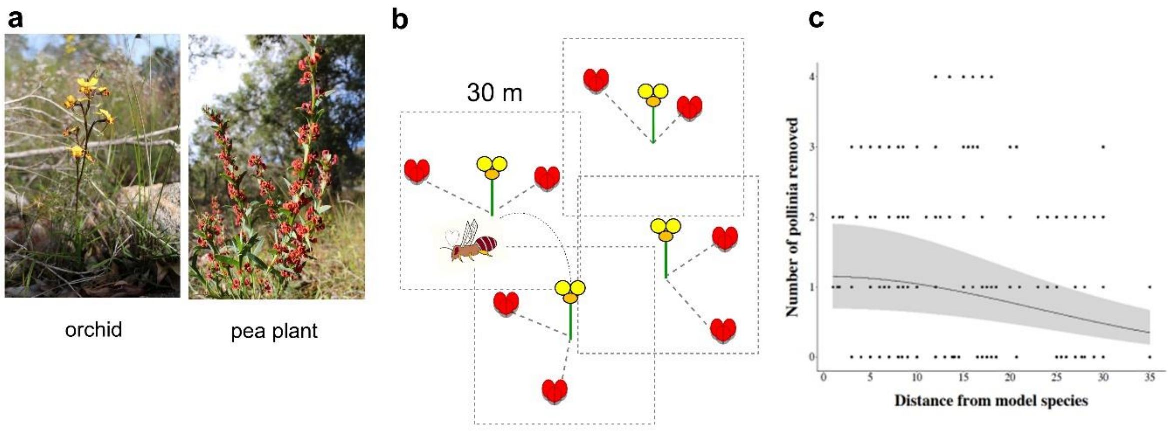


**Fig. S2.** Orchid pollination success relative to distance from model pea plants. (a-b) The effect of relative distance between an individual orchid (yellow flower) and the surrounding pea models (red flower) was quantified within 30 x 30 m quadrat centered on the orchid plant (N = 122 orchids across 5 populations). (c) The number of pollinaria removed from the orchid flowers decreased significantly with 9 the square of orchids' distance from the pea model ($\chi^2$ =17.09, $p < 0.001$). Specifically, pollination success was compromised when the relative 0.002) distance between orchid and pea was greater than 15 m ($\chi^2$ =9.49, p = 0.002).

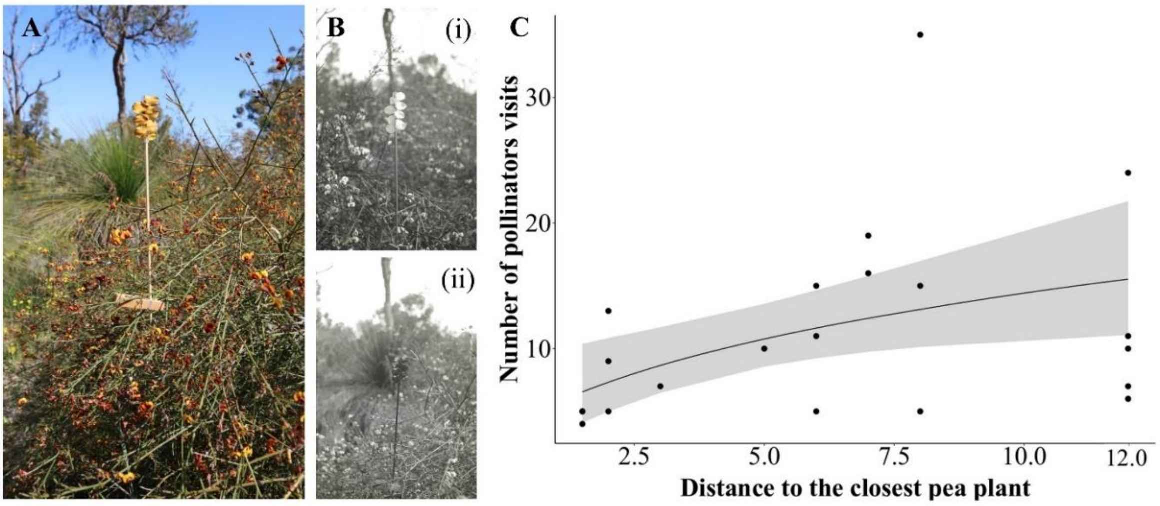


**Fig. S3. Experimental manipulation of *Daviesia divaricata* floral display using a 'UV flag'. (A)** A 'UV flag' was made by attaching the outer petals of *Diuris magnifica* on a wood stick to exaggerate the UV reflectance floral display of the pea *Daviesia divaricata*. **(B)** Ultraviolet photography of the 'UV flag' (i) untreated and (ii) treated with the ultraviolet screening. **(C)** The visiting rate of treated *Daviesia* was not influenced by their distance to the closest pea plant ($\chi^2$= 1.02, p = 0.313). However, the visiting rate of *Daviesia* when its UV reflectance was exaggerated by the untreated UV flag was affected by the proximity to other *Daviesia* plants ($\chi^2$= 6.29, p = 0.012) and increased logarithmically with distance from the closest pea plants.

**Supplementary Information Text**

**Methods S1**

**Testing the effect of the ultraviolet reflectance screening solution on pollinator visits.** The effect of the UV reflectance screening solution (Kinesys, Canada) on the number of *Trichocolletes* bee visits to *Diuris* orchids was tested using choice experiments conducted during the foraging peak of pollinators (11.00 -14.00) over two days. Two picked inflorescences with identical number of flowers were presented in proximity to model plants according to rotating arrays methodology (Scaccabarozzi et al., 2020a). One of the two inflorescences was treated by applying the screening solution at the base of the corolla, whilst the other one was used as control (non-treated). Number of bees approaching the inflorescences with treated and untreated flowers was recorded for 16 trials and subsequently tested by a Generalized Linear Model with a Bernoulli distribution of the response variable. We found no difference between the number of *Trichocolletes* bees visiting treated (n = 29) and non-treated orchid flowers (n = 31; $\chi^2 = 1.11$, $p = 0.291$). We concluded that the screening solution had no effect in attracting to or repelling the pollinators from orchid flowers and thus, suitable for the UV manipulation experiments conducted in the current study.

**Control experiment for olfactory cues.** The ability of *Trichocolletes* bees to detect orchid flowers by olfactory cues was also tested by obscuring flowers using a black PTE cylinder as per Phillips et al. (2014). Choice experiments were conducted during the foraging peak of pollinators (11.00 -14.00) over two days. Two picked inflorescences with identical number of flowers were presented in proximity to model plants according to a rotating arrays methodology[1]. One of the inflorescences was obscured using a black cylinder, 20 cm high but open at the top and raised 10 cm off the ground, allowing access to the hidden flowers. Every 15 min-trial their position was changed relative the model plants, maintaining the same distance between the two inflorescences (30 cm). The number of bees approaching the inflorescence with the flowers hidden by the dark cylinder and the inflorescence with visible flowers was recorded for 24 trials. Number of bees approaching hidden and not-hidden orchid flowers was tested by a Generalized Linear Model with a Bernoulli distribution of the response variable. Bees attracted by hidden and not-hidden orchid differed significantly ($\chi^2 = 152.49$, $p < 0.001$). Specifically, we found that a total of 55 *Trichocolletes* bees approached the hidden orchid flowers, whilst none of them approached the flowers hidden by the dark cylinder. As expected, *Diuris* attracts *Trichocolletes* bees by visual cues and not by olfactory cues.

**Supplementary Information Text**

**Methods S2**

**Pilot for testing the generality of UV reflectance as a long-range signal**

We investigated whether the long-range UV signalling observed in the orchid *Diuris brumalis* is a phenomenon found in other plant species, which are taxonomically distant to orchids. The generality of long-range UV signalling was tested by manipulating the floral display of the pea plant, *Daviesia divaricata*, in a pilot field experiment.

Like other *Daviesia* spp., *D. divaricata* is pollinated by *Trichocolletes* bees that also pollinate the co-flowering mimicking orchid species *Diuris magnifica* (Scaccabarozzi et al., 2020b). Unlike other congeneric species, however, this pea species is also pollinated by a more generalist range of pollinators including native bees and introduced honeybees (Scaccabarozzi et al., 2020c). To test the attraction rate of these pollinators to *D. divaricata* plants, we conducted a total of 42 trials of five-minute observation of floral visitors in 21 pea plants in a pea plant patch with intense bee activity across four days from 21 to 24 September 2021. We used a 'UV flag' made by attaching the outer petals of the orchid *D. magnifica* on a stick to exaggerate the UV reflectance of *Daviesia* floral display (i.e. untreated). This manipulation was possible because the colouration of the standard petal in the pea *Davesia divaricata* matches that of the outer petals in the orchid *Diuris magnifica* (i.e. the colour falls in the UV sector of the bee visual model hexagon) (Scaccabarozzi et al., 2020b), making the two species undistinguishable by bees.

In half of the trials (N = 21), the 'UV flag' was treated with the same clear ultraviolet screening solution used in our previous experiments to eliminate ultraviolet reflectance (i.e. treated; Fig. S3A, B). Observational trials using treated and untreated 'flags' were conducted one after the other to minimise temporal shifts in observation. Only bees flying from further away and then approaching the flowers of a single pea plant were recorded. Bees were identified and classified per pollinator taxon by visual assessment in: *Trichocolletes*, *Apis mellifera*, *Leioproctus*, *Megachilidae* and *Halictidae*. Trials were conducted by selecting pea plants for observation and by recording the distance to the closest pea plant. We selected pea plants that were not surrounded by other pea plants to avoid the influence of other magnet species into pollinator attraction. We found that the long-range ultraviolet signalling increases the detectability and visitation rates of the *Daviesia* pea (Fig. S3C) up to 12 meters. This finding indicates that the phenomenon is likely to be widespread across plant taxa. For example, a range of pea plants in south-west Australia have been shown to display a similar colour pattern to *Daviesia* (i.e. 'egg and bacon' peas) (Scaccabarozzi et al., 2020c) and are likely to use a long-range UV signalling as well.

Data were analyzed by a GLM with the number of bee visits (all species combined) as response variable and the distance from the closest pea plants as covariate. The dependency from the distance was assessed considering the interaction with the treatment and the shape of the predictors was determined by using the AICc criterion for model comparison. The distribution exploited was a negative binomial to account for the overdispersion occurring in the Poisson model (overdispersion parameter: 4.09). The overdispersion parameter of the negative binomial model was 1.25 and its affordability for the goodness of the model was assessed through a simulation study to evaluate whether the number of zeros from the model was included in tolerability range. Results were suitable to consider the model reliable. All the analyses were conducted by R Studio (Version 1.4.1106). Package exploited were MASS, MuMIn, plyr and ggplot2.

**Supplementary Information Text**

**Methods S3**

**Conversion of photos of *Diuris* sp. and *Daviesia* sp. in false colour photography**

False colour photography in ‘bee view’ format was used to reveal the overall colour pattern perceived by bees in treated (i.e., application of the UV screen solution) and untreated outer petals of *Diuris* flower and untreated *Daviesia* flower. Each flower was photographed in colour (i.e., human view) and UV using a modified Panasonic BMC-G3 camera with a UV-transmitting Ultra-achromatic-Takumar 20 mm F/4 lens made of fused quartz. The white balance was set separately for the colour and UV-photography using a white Teflon disc as a control under same light conditions. The images, taken from the same position and within 10 sec intervening time, were converted into false colour images that comprise the UV, blue and green wavelengths perceived by bees. Image assemblage in bee view was obtained by splitting the colour and the UV-photo into the three camera colour channels, namely blue, green and red. The red channel was discarded from the colour photo and green and red channels were discarded from the UV-photo as per Lunau et al. (2021). In the false colour photos UV wavelengths are represented as blue, blue as green and green as red wavelengths as a standard method for colour image translation to represent bee vision. In bee view, *Diuris* petals treated with the UV filtering solution exhibited a different coloration (orange, characterised by green wavelengths) compared to untreated petals (purple, characterized by UV wavelengths and green wavelengths).

**Table S1.** List of the five population sites of *Diuris brumalis* surveyed.

| Population number | Site | Latitude, longitude |
|---|---|---|
| 1 | Lesmurdie - Canning Rd | 32°01'45.6'' °S, 116°06'01.3'' °E |
| 2 | Lesmurdie - Canning Rd | 32°01'41.9'' °S, 116°05'41.7'' °E |
| 3 | Lesmurdie - Canning Rd | 32°01'43.8'' °S, 116°05'12.1'' °E |
| 4 | Lesmurdie - Canning Rd | 32°01'43.2'' °S, 116°04'51.9''° E |
| 5 | Lesmurdie - Canning Rd | 32°01'39.8'' °S, 116°04'45.3'' °E |

**References**

Lunau, K., Scaccabarozzi, D., Willing, L., & Dixon, K.W. (2021). A bee's eye view of remarkable floral colour patterns in the Southwest Australian biodiversity hotspot revealed by false colour photography. *Annals of Botany*, *128*, 821–834.

Phillips, R. D., Scaccabarozzi, D., Retter, B. A., Hayes, C., Brown, G. R., Dixon, K. W. & Peakall, R. (2014). Caught in the act: pollination of sexually deceptive trap-flowers by fungus gnats in *Pterostylis* (Orchidaceae). *Annals of Botany, 113,* 629-641.

Scaccabarozzi, D., Galimberti, A., Dixon, K. W., and Cozzolino, S. (2020a). Rotating Arrays of Orchid Flowers: A simple and effective method for studying pollination in food deceptive plants. *Diversity*, *12*, 286.

Scaccabarozzi, D., Guzzetti, L., Phillips, R.D., Milne, L., Tommasi, N., Cozzolino, S., and Dixon K.W. (2020b). Ecological factors driving pollination success in an orchid that mimics a range of Fabaceae. *Botanical Journal of the Linnean Society, 194*, 253-269.

Scaccabarozzi, D., Dixon, K. W., Tomlinson, S., Milne, L., Bohman, B., Phillips, R. D. & Cozzolino, S. (2020c). Pronounced differences in visitation by potential pollinators to co-occurring species of Fabaceae in the Southwest Australian biodiversity hotspot. *Botanical Journal of the Linnean Society*, *194*, 308-325.

**Legends for Data S1 to S6**

**Data S1. Morphological measurements of *Diuris* and *Daviesia* floral components and UV salient signal ratio calculation.** Flower measurements (a, b, r) taken on *Diuris* for calculating the area of the geometric figures (ellipse and circle) that approximate the area of the flower components (see Fig. S1). a, b: major and minor axis of the outer petal; r: distance between the pollinaria centre and the top edge of the dorsal sepal. The ellipse approximates the outer petal area that has been duplicated for estimating the total area of the external flower component, formed by the two outer petals. Flower measurements (a1, b1, a2, b2) taken on *Daviesia* for calculating the area of the geometric figures (ellipse 1 and 2) that approximate the area of the flower components (see Fig. S1). a1, b1: major and minor axis of the wing petals comprising the keel edge. a2, b2: major and minor axis of the two standard petals. Calculations of UV salient signal ratio with total surface of flowers (comprising all flower components) and total surface of flowers reflecting more than 10% in UV, according to cut value by Dyer (1996) for Australian plants. *between 300 and 400 nm (full spectra reflectance curve above 10%); [1]outer petals and dorsal sepal [2]: standard petal; *Diuris* dorsal sepal: estimated as semi-circle area of the flower central component in Fig. S1. *Daviesia* standard petals: estimated by subtracting the area of Ellipse 1 to the area of Ellipse 2 in Fig. S1.

**Data source S2. Spectral measurements of the floral components of *Diuris brumalis* and *Daviesia decurrens.*** Means and standard deviation of colour reflectance for *Diuris* and *Daviesia*. Means are based on colour measurements across flower components of six individual plants.

**Data source S3. Testing the effect of the UV screening spray on the attraction of *Trichocolletes* bees.** Number of bees approaching *Diuris* inflorescences with UV treated and untreated flowers.

**Data source S4. Testing *Trichocolletes* bee attraction by olfactory cues of *Diuris*.** Number of bees approaching hidden and visible orchid flowers.

**Data source S5**. **Model-mimic distance experiment.** Testing whether *Diuris* pollination success varies depending on the distance to the model pea plants.

**Data source S6. In and Out experiment.** First field UV manipulation experiment, testing that UV reflectance enhances orchid pollination success when out from peas patch.

**Data source S7. Distance from orchids to model plants as continuous variable.** Second field UV manipulation experiment, testing that by displaying an exaggerated version of *Daviesia*'s attractive UV reflectance, *Diuris* benefits from pollinators that mistake it for the rewarding model from afar.
